# Background reduction in STED-FCS using coherent-hybrid STED

**DOI:** 10.1101/2020.01.06.895243

**Authors:** Aurélien Barbotin, Iztok Urbančič, Silvia Galiani, Christian Eggeling, Martin Booth

## Abstract

Fluorescence correlation spectroscopy (FCS) is a valuable tool to study the molecular dynamics of living cells. When used together with a super-resolution stimulated emission depletion (STED) microscope, STED-FCS can measure diffusion processes at the nanoscale in living cells. In twodimensional (2D) systems like the cellular plasma membrane, a ring-shaped depletion focus is most commonly used to increase the lateral resolution, leading to more than 25-fold decrease in the observation volumee, reaching the relevant scale of supramolecular arrangements. However, STED-FCS faces severe limitations when measuring diffusion in three dimensions (3D), largely due to the spurious background contributions from undepleted areas of the excitation focus that reduce the signal quality and ultimately limit the resolution. In this paper, we investigate how different STED confinement modes can mitigate this issue. By simulations as well as experiments with fluorescent probes in solution and in cells, we demonstrate that the coherent-hybrid (CH) depletion pattern reduces background most efficiently and thus provides superior signal quality under comparable reduction of the observation volume. Featuring also the highest robustness to common optical aberrations, CH-STED can be considered the method of choice for reliable STED-FCS based investigations of 3D diffusion on the sub-diffraction scale.

## 1 Introduction

Fluorescence correlation spectroscopy (FCS) is a technique that allows observation of molecular diffusion at the scale of a couple of hundreds of nanometers [1], [2]. Traditional FCS implementation makes use of confocal optics, and as such is limited by the diffraction limit to lengthscales of approximately 200 nm. This can be a problem when studying nanoscale organisation, for instance in the plasma membrane. Fortunately, the resolution of FCS microscopes can be increased by means of stimulated emission depletion (STED) (STED-FCS [3]) to access the diffusion information on the sub-diffraction scale. In STED microscopy, the excitation laser beam is overlaid with a high intensity depletion laser beam that exhibits a central intensity minimum, inducing stimulated emission in the areas of its high intensity and thus reducing the effective observation volume to the very center of the excitation focus. The depletion pattern is thus a key component of a STED system. The most commonly used depletion pattern is a ring-shaped focus, which constrains the lateral resolution but leaves unchanged the axial resolution (2D STED). This depletion pattern has been extensively used to study two-dimensional systems like cellular membranes [4]–[8], but faces severe limitations when studying three-dimensional diffusion, due to varying axial cross-sections and high background originating from undepleted areas [5], [9] that leads to an increase in the apparent number of molecules in the observation volume [5], [10]. Alternatively, a bottle-shaped depletion beam can be used to essentially constrain the axial resolution (z-STED), but is more sensitive to optical aberrations [11], [12], which can be mitigated with adaptive optics [13]. However, even aberration-free z-STED-FCS exhibits spurious contributions from undepleted areas that damp the amplitude of the autocorrelation function, biases measurements of the numbers of molecules and reduces the signal-to-noise ratio [3], [5]. A combination of both 2D and z-STED depletion patterns (3D STED) has also been used in STED-FCS [5], [14] but did not significantly reduce background contributions. More recently, contrast in STED imaging has been increased using a superposition of two mutually coherent ring-shaped foci created by a bi-vortex phase mask [13], named coherent-hybrid (CH) STED. Anticipating signal-to-noise improvements also in STED-FCS, we here investigate the origins of background in STED-FCS experiments performed with common STED confinement modes (2D-, z-, and 3D-STED) as well as with CH-STED. We characterised the performance and sensitivity to optical aberrations of each of these confinement modes in STED-FCS, and showcased their use in biological specimens.

## 2 Methods

### 2.1 Microscopes

Experiments were performed using a custom STED microscope built around a RESOLFT microscope by Abberior Instruments equipped with an oil immersion objective lens (Olympus UPLSAPO, 100×/1.4 oil), as described in previous publications [13], and as sketched in Figure 1.

**Figure 1:**
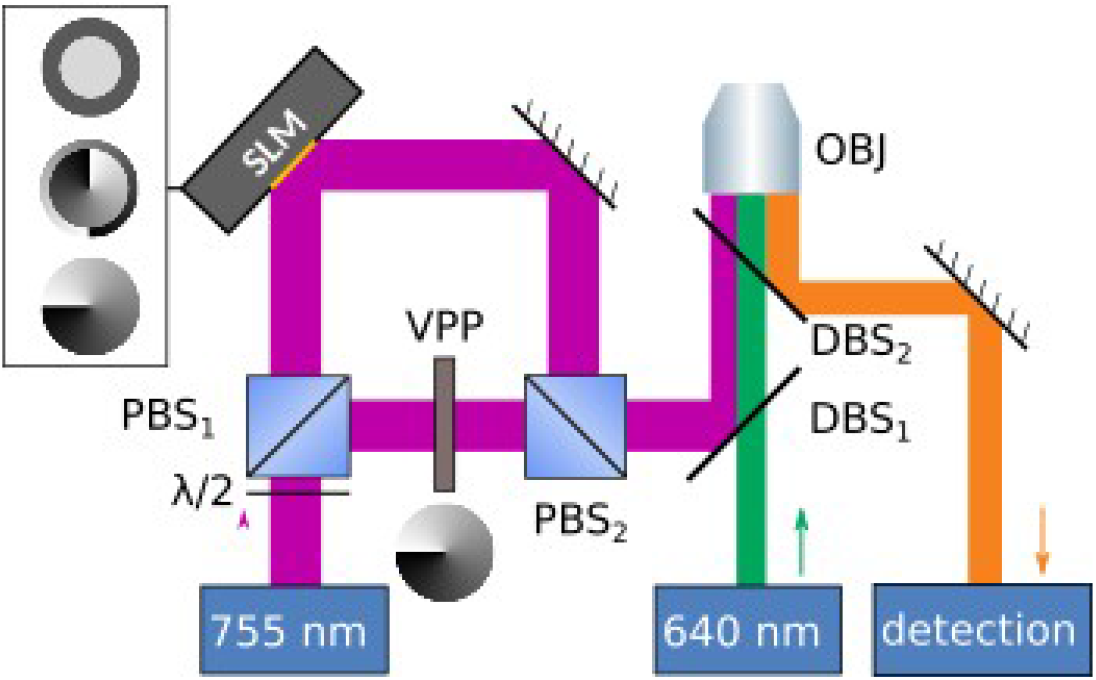
Sketch of the microscope. Excitation (green) and depletion (magenta) lasers are focussed by an oil immersion objective (OBJ). The depletion laser beam is split in two orthogonal polarisations and later recombined using a polarisation beam splitter (PBS 1 and 2). One component is modulated in phase by the SLM (gray box) that generates different phase patterns (insert), while the other is modulated by a vortex phase plate (VPP). Excitation, depletion and detection (orange) beam paths are recombined using dichroic beamsplitters (DBS 1 and 2).

The depletion STED laser (Spectra-Physics Mai Tai, pulse-stretched by a 40-cm glass rod and a 100-m single-mode fibre) was pulsing at a frequency of 80 MHz at a wavelength of 755 nm. The STED laser beam was separated in two arms using a polarisation beam splitter, with the amount of light going in each arm being controlled by rotating the plane of the incident linear polarisation using a λ/2 phase plate. In the first arm, a spatial light modulator (SLM, Hamamatsu LCOS X10468-02) was used to generate 2D-, z- and CH-STED patterns. In the second arm, a vortex phase plate (VPP-1a, RPC Photonics, Rochester, NY) was used to create a 2D depletion pattern, which was overlaid with a z-STED pattern generated by the SLM to create 3D STED. System aberrations in the depletion arm including the SLM were removed by scanning a sample of scattering gold beads, using the sensorless method and using image standard deviation as an image quality metric. A 640 nm laser pulsing at a frequency of 80 MHz was used for excitation, at powers ranging from 4 to 17 μW measured in the back focal plane of the objective.

### 2.2 Depletion patterns

We investigated the performances of in total 4 depletion patterns: 2D-, z-, CH-, and 3D STED (Figure 2). 2D-, z- and CH-STED patterns were generated with the SLM, while the 3D STED pattern was created as an overlay of an SLM-generated z-STED pattern and a phase-plate generated 2D STED pattern; if not stated otherwise, 80% of the STED laser power was in the z-STED arm and 20% in the 2D STED. The inner radius of the CH-STED mask can be changed to modify the shape of the corresponding depletion pattern. We initially set the value of this parameter to 0.85 (85% of the pupil radius size), as was used in a previous implementation [15]. The influence of the CH-STED radius parameter is discussed in section 4.1.

**Figure 2:**
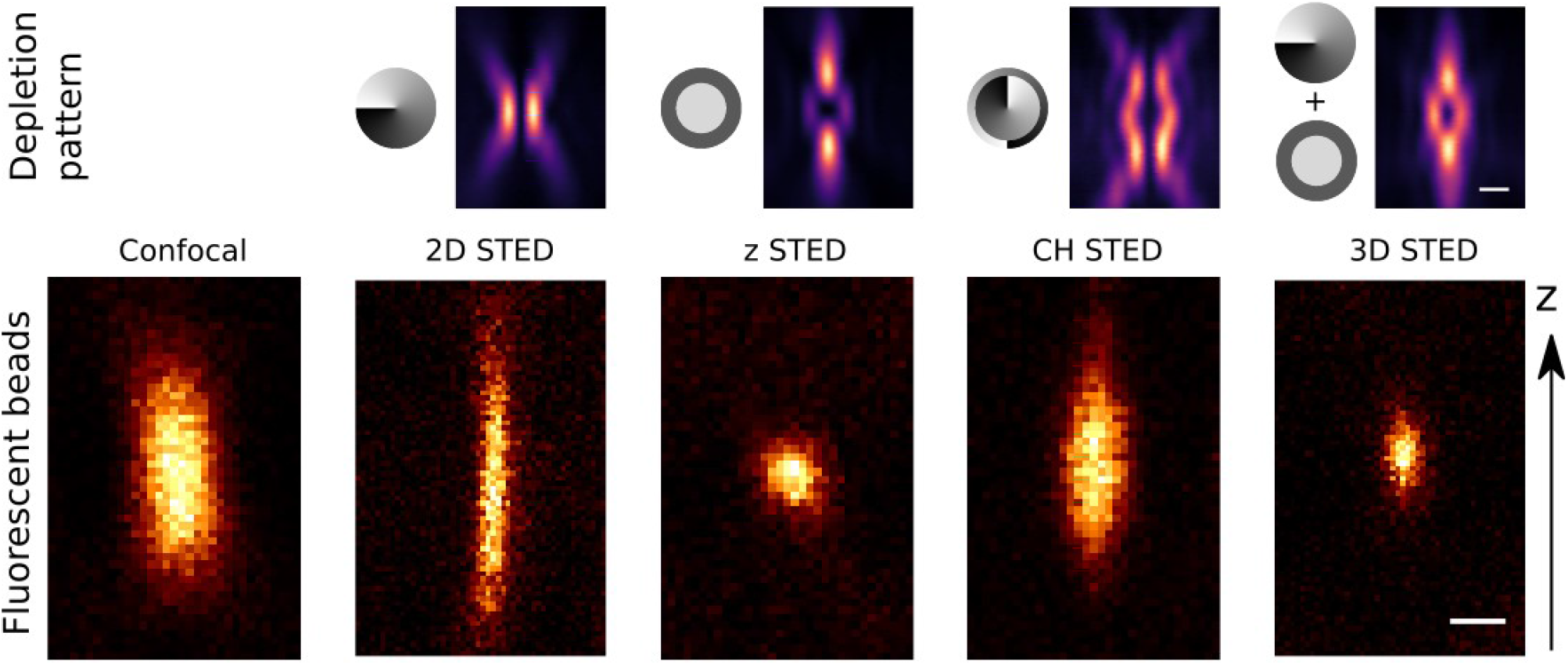
Phase masks used for STED-FCS. Top: Schematics of the phase patterns of the masks (left, black to white: phase shift from 0 to 2π) used to generate the corresponding STED depletion patterns (right, axial cross-sections of back-reflected images of gold beads, scalebar: 500 nm). Bottom: experimental images of 40 nm fluorescent beads, acquired in confocal (left) and with the different STED confinement modes, at a STED laser power of 55 mW. Scalebar: 200 nm.

### 2.3 Samples

#### Dyes solution

Freely diffusing dyes in solution were prepared by diluting Abberior Star Red dyes (Abberior, Germany) to a concentration of 50 nM in a 1:1 water:glycerol solution. Glycerol was used to slow down diffusion speeds to facilitate the analysis of FCS data.

#### Fluorescent beads

Slides of immobilised 40 nm far-red fluorescent nanoparticules used for STED imaging were purchased from Abberior Instruments (Germany).

#### Supported lipid bilayers

Supported lipid bilayers were prepared as described previously [16]. The cover slips were cleaned with piranha acid (3:1 sulphuric acid and hydrogen peroxide) and stored in water for no more than 2 weeks before experiment. 25 μL of 1 mg/mL POPC (1-palmitoyl-2-oleoyl-sn-glycero-3-phosphocholine; Avanti Polar Lipids, AL, US) lipid solution in chloroform/methanol with 0.01 mol% of fluorescent Abberior Star Red-labelled phosphatidylethanolamine (PE; Abberior) were spin-coated onto a clean dry coverslip at 3,200 rpm for 30 seconds. The lipid film was rehydrated with SLB buffer (10 mM HEPES and 150 mM NaCl pH 7.4) and washed several times to remove non-planar lipid structures.

#### Cells

Cells were prepared using the same protocol as in [11]. Human fibroblasts (GM5756T, Moser, Baltimore, USA) were maintained in a culture medium consisting of DMEM with 4500 mg glucose/L, 110 mg sodium pyruvate/L supplemented with 10% fetal calf serum, L-glutamine (2 mM) and penicillin-streptomycin (1%). The cells were cultured at 37 °C/5% CO_2_. Cells were grown in a 35 mm imaging dish with a glass coverslip bottom (ibidi GmbH, Germany), and transfected with a plasmid expressing a fusion protein of GFP and SNAP-tag using Lipofectamine 3000 transfection reagent (Invitrogene, Carlsbad, USA). 24 hours after transfection, the cells were incubated together with SNAP-Cell 647-SiR (New England Biolabs (UK) Ltd., Hitchin, UK) and washed twice in culture medium after 40 min incubation, with a waiting time between washings of 20 min. Finally the culture medium was substituted with L-15 medium (Sigma-Aldrich, Dorset, UK) and each sample was visualized at 37°C for no longer than 1 hour.

### 2.4 FCS

FCS curves were either obtained directly from a correlator card (Flex02-08D, correlator.com) or by acquiring fluorescence intensity timetraces with a frequency comprised between 0.25 and 1 MHz that were correlated offline using the python package multipletau [17]. Acquisition times were set to 10 seconds.

FCS parameters were obtained by fitting FCS curves with standard diffusion model, assuming Gaussian-shaped observation volumes [18]:

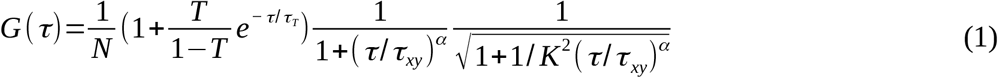

Where *N* refers to the average number of molecules in the effective observation volume, *T* is the average triplet amplitude, *τ_T_* is the triplet correlation time, *τ_xy_* is the average lateral transit time, *α* is a factor characterising deviation (values different from 1) from the Gaussian shape of the observation volume or anomalous diffusion, and *K* is the aspect ratio of the observation volume, defined as *K* = *ω_z_*/*ω_xy_*, and *ω_xy_* and *ω_z_* are respectively the lateral and axial 1/e^2^ radii. The lateral transit time *τ_xy_* and *ω_xy_* size are related to the diffusion coefficient *D* [1], [2], [18]:

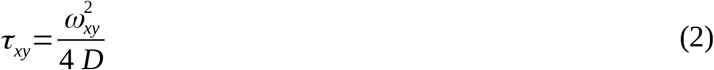

Triplet correlation times *τ_T_* were determined from confocal recordings, and set to the constant value of 12 μs for Abberior Star Red in solution and of 5 μs otherwise. A general procedure for STED-FCS data analysis can be found in reference [6]. The shape factor *α* was also determined from confocal recordings, and was set to 1 (describing a Gaussian profile and free diffusion) in every sample except for cells, where *α* was set to 0.8 to account for anomalous subdiffusion caused by the crowded environment in the cell cytoplasm [6], [19]. Since typical molecular brightness of conventional fluorophores is not high enough to independently determine the lateral transit time *τ_xy_* and the aspect ratio *K* [3], [5], we calibrated the variations of aspect ratio with lateral size for each STED confinement mode, and fitted observation volumes with a prescribed shape, as described in [13] (see supplement S1).

### 2.5 Adaptive optics

The employed depletion patterns show different sensitivity to optical aberrations [11], [20], [21]. To ensure optimal performance and thus fair comparison of the confinement modes, we first corrected sample-induced optical aberrations, as well as residual system-induced aberrations, using the sensorless adaptive optics approach described in [13]. While measuring z-STED FCS (the most aberration-prone mode), we corrected low-order Zernike modes (modes 5 to 11, following the convention defined by Noll [22]) in each sample by minimising the average number of molecules in the observation volume. The correction optimises the shape of the depletion beam for all confinement modes, but can result in their slight misalignment [23]. Hence, once the correction was determined using z-STED-FCS, coalignment between excitation and depletion pattern of interest (2D or CH) was ensured by optimising tip and tilt, again using the average number of molecules in the observation volume as a quality metric. During aberration correction procedures, STED laser power was set to 16 mW. Aberration amplitudes were measured in radians root mean square (rad rms, see section 4.3 and supplementary material) and measured at 755 nm.

### 2.6 Simulations

The intensity distributions of the excitation and depletion lasers were calculated using the vectorial diffraction theory, as described for instance in [23]. Depletion patterns were simulated at a wavelength of 755 nm, and excitation was calculated at 640 nm, both wavelengths used in our system. The detection profile was defined as a convolution of the excitation profile with a pinhole with a size of 1 Airy unit. Pixel size was set to 10 nm. To minimise computing time, only 2-dimensional (xz) profiles were calculated. Integration was performed in cylindrical coordinates, using the invariance of the calculated foci along the azimuthal coordinate to calculate threedimensional integrals from two-dimensional simulation data.

### 2.7 Estimation of background contributions with SLBs

In section 3.2, FCS measurements were performed on SLBs, at different axial positions of the excitation and STED focus with respect to the membrane. At each position, 3 FCS curves were recorded with an acquisition time of 10 s. Resulting curves were fitted with the model from equation 1 to extract the average number of molecules in the observation surface and the average transit time. The average photon count was also recorded. From these three quantities, we could estimate background levels as follows. Undepleted background increases the apparent number of molecules in the observation surface [5], [9], [10]:

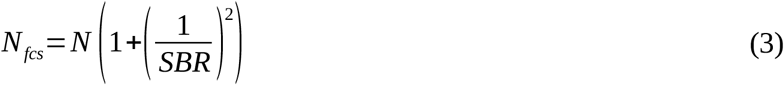

where *N_fcs_* is the number of molecules in the observation volume estimated with FCS, *N* is the actual number of molecules in the observation volume and SBR is the signal-to-background ratio, defined as *SBR* = 〈*F*〉/〈*F* + *B*〉, where *F* is the correlating fluorescence, *B* is the uncorrelated background fluorescence and 〈.〉 designates the time-averaging operator.

At a depth z, *N_fcs_* can be directly obtained from the amplitude of FCS curves. The actual number of molecules N in the observation surface is a function of the concentration of fluorescent molecules per surface unit *c* and of the size of the observation surface *S*:

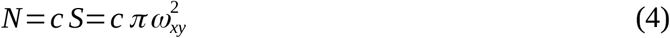

The concentration *c* is unknown, but its determination is not necessary: assuming that there is no undepleted background at depth 0, when the SLB is in the focal plane of the objective, we have *N*_0_ = *N*_*fcs* 0_, where *N_0_* is the actual number of molecules in the observation surface at depth 0, and *N*_*fcs*0_ is the average number of molecules in the observation surface at depth 0, measured by FCS. Equation 4 at any depth z can then be divided by the values measured at depth 0:

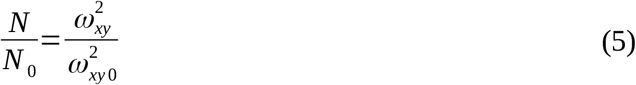

Where *ω_xy,0_* is the size of the observation surface at depth 0, which was determined as the plane of measurements with the highest photon counts. At a depth z, the number of molecules can be calculated from equations 2 and 5 as:

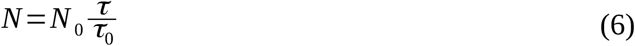

Finally, at each depth, we can calculate the SBR:

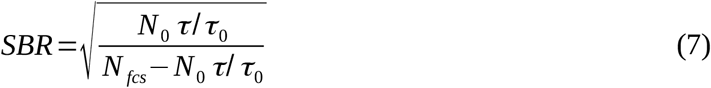

This quantity assumes that *N_fcs_* is systematically larger than *N*, which is always true according to the theory (equation 3). However, in situations where very low background is present, statistical variations in the measurements of *N* and τ can lead *N_fcs_* to be slightly smaller than *N*. In this case, we considered *N_fcs_* and *N* equal and set the SBR to an infinite value. Determination of the absolute size of the observation surface results directly from equation 2:

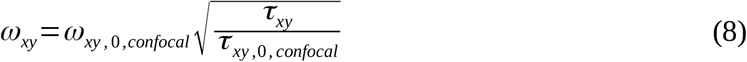

Where *ω*_*xy*,0,confocal_ is the lateral size of the confocal spot in the focal plane, measured to be 102 nm (corresponding to a full width at half maximum of 240 nm) by imaging a sample of fluorescent beads, and τ_xy,0,confocal_ is the lateral transit time as determined in SLBs with confocal FCS. Knowing the lateral size of the observation volume and the average photon counts at each depth, we could reconstruct the Gaussian intensity profile of each focus using:

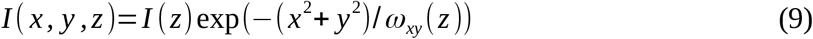

Where *I* refers to the average photon counts. Knowing the SBR at each depth from equation 7, we calculated at each depth the fraction of photons contributing to the signal and the fraction contributing to background:

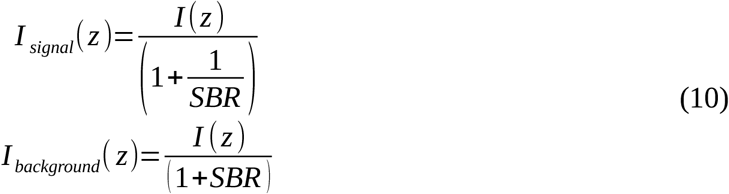

As an alternative measure of the noise in experimental FCS curves, we calculated the root-meansquare values of the fitting residuals up to a lag time of 50 μs (smaller than transit times), normalised by the amplitude, as applied before [24].

## 3 Origin of background in 3D STED-FCS

### 3.1 Simulations

Recent work showed that the relatively poor signal-to-background ratio (SBR) obtained with 2D STED-FCS when studying 3D diffusion was due to low-intensity contributions originating from undepleted areas of the excitation focus [5], [9]. To extend this study to different STED confinement modes, we first obtained better insight into spatial origins of signal and background by simulating the excitation and depletion foci of our system.

In the resulting effective intensity distributions in confocal and different STED confinement modes (Figure 3), low- and high-intensity areas, which yield respectively weakly and strongly correlating contributions, can be considered as regions giving rise to background and signal, respectively. In practice, we used a threshold defined as 1/e^2^ of the maximum intensity (black dashed contours in Figure 3), as previously in STED-FCS [9]. Because intensity distributions are symmetric with respect to the optical axis, they are conveniently represented in cylindrical coordinates (Figure 3(a), top). In this representation, however, not all pixels contribute equally to the overall intensity, as the value of the integral along the azimuthal axis increases with distance from the optical axis. A more informative representation of signal and background contributions is thus obtained by integrating the intensity distribution along the azimuthal axis (Figure 3(a), bottom). In this representation, the 1/e^2^ threshold in plain intensity distribution was used to separate pixels contributing to signal and background, which were plotted with different colorscales for clarity.

**Figure 3:**
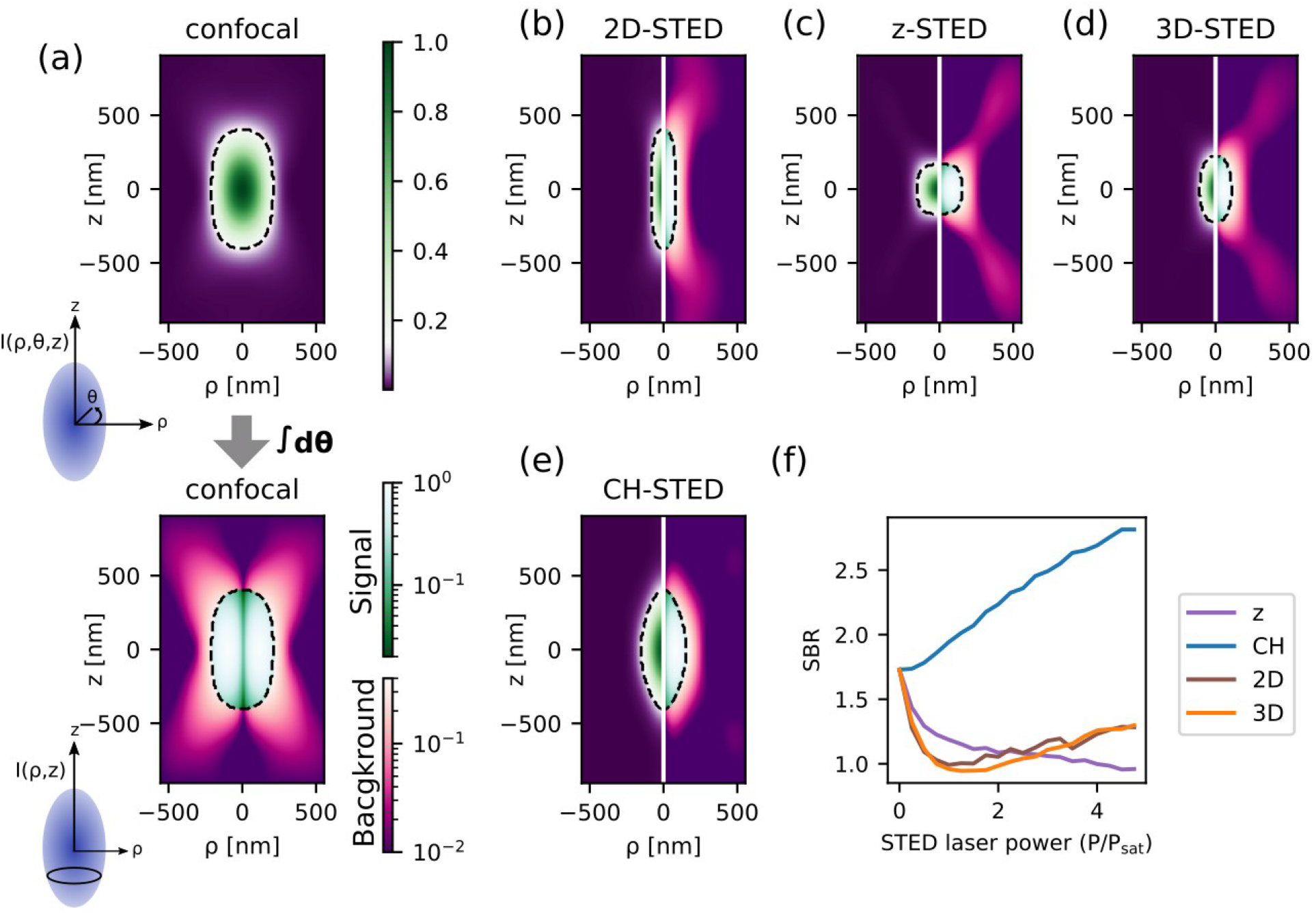
Simulation of spatial distribution of signal and background in STED-FCS. (a) Visualisation of spatially-varying signal and background contributions in a confocal observation spot, for the axial cross-section of the intensity distribution (top) and integrated along the azimuthal axis (bottom) as sketched on the left. Dotted black lines: contours at 1/e^2^ of the maximum intensity. (b)-(e): Visualisations of spatially-varying signal and background contributions for 2D-STED (b), z-STED (c), 3D-STED (d) and CH-STED (e), using the same colormap as in (a). Plain intensity distribution is displayed on the left, and intensity integrated along the azimuthal axis is displayed on the right. (f): Signal-to-background ratio (SBR) integrated across all three dimensions varying with STED laser power (normalised to the saturation power P_sat_, defined as the STED laser power leading to a 2-fold lateral resolution improvement in 2D STED), for different STED confinement modes as indicated in the legend.

Our simulations showed that background values in both confocal and 2D STED were significant at large distance from the focal plane (higher than 300 nm, Figure 3(a)-(b)). In z-STED (Figure 3(c)), significant background contributions were similarly caused by undepleted out-of-focus side lobes, which were only slightly reduced by 3D-STED (Figure 3(d)). CH-STED (Figure 3(e)), on the contrary, reduced background values in all areas of the observation focus, thanks to a good overlap between excitation and depletion foci.

By integrating the spatial contributions to signal and background, we obtained an estimate for an overall SBR (Figure 3(f)). Our simulations showed that CH-STED suppressed background most effectively, resulting in considerably higher SBR compared to other confinement modes, and thus promising superior signal quality of CH-STED in actual FCS measurements.

### 3.2 Experimental measurement of background

We next validated our model experimentally using supported lipid bilayers (SLBs). SLBs exhibit 2-dimensional diffusion, and by measuring with (STED)-FCS at different axial positions of the focus with respect to the membrane plane (Figure 4(a)), we determined the out-of-focus uncorrelated background contributions as would arise in the case of 3D diffusion. For distances to the focal plane ranging from 0 to 1100 nm, we measured both the average number of molecules in the observation surface and the average transit time, which is proportional to the size of the observation surface (equation 2). In the absence of undepleted background light, the variations in average number of molecules in the observation area and in the size of observation surface should be strictly proportional. In presence of background light, however, the apparent number of molecules in the observation surface (*N_fcs_*) increases faster than the size of the observation surface due to a damping of the correlation amplitude. The difference between expected and actual number of molecules in the observation surface thus allowed calculating the contributions of signal and background as described in the methods section (section 2.7).

**Figure 4:**
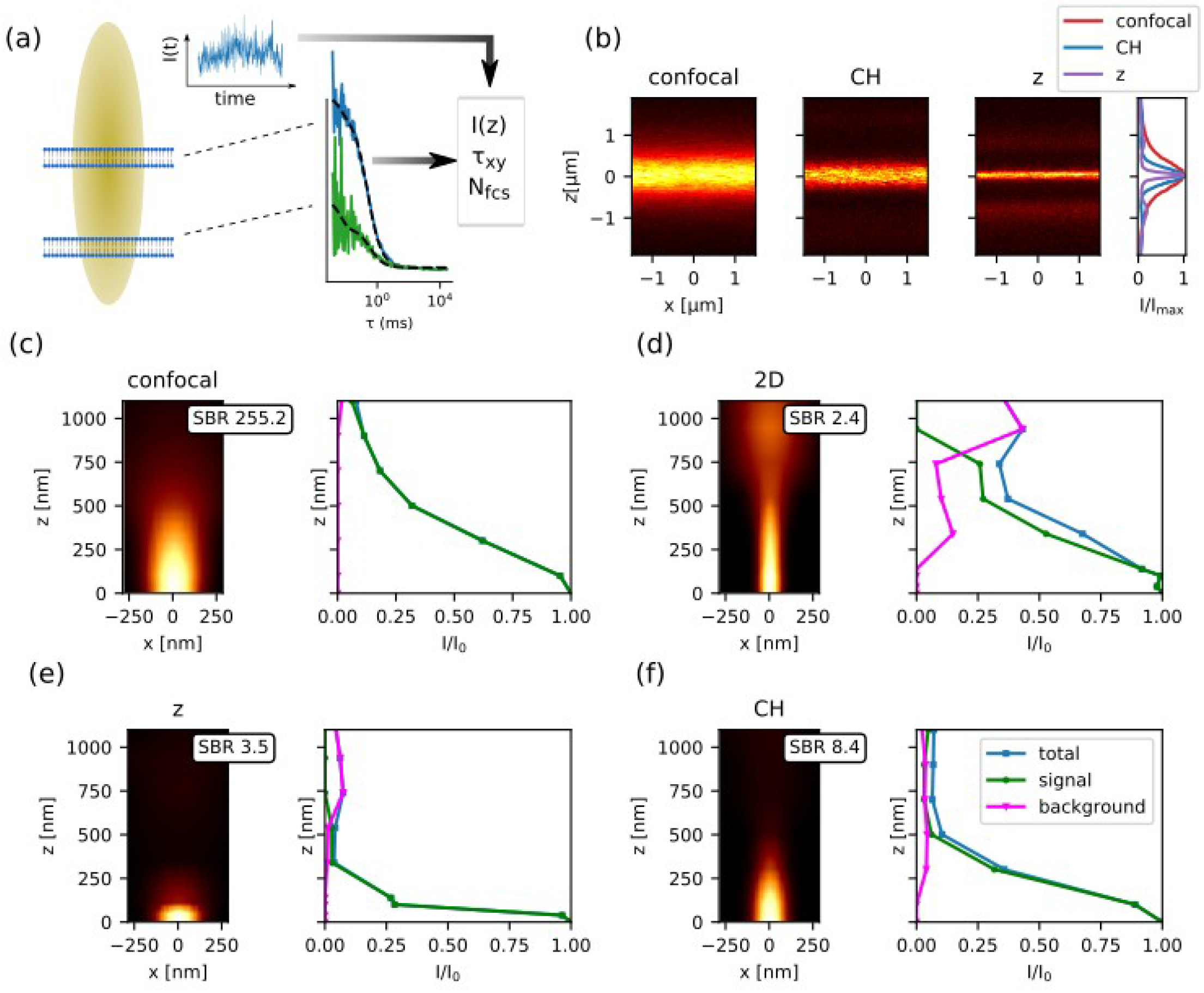
Experimental determination of background in STED-FCS using supporting lipid bilayers (SLBs). (a) Principle of the experiment: FCS measurements on planar SLBs at various axial positions of the focus (z) were used to probe the intensity (I), size of the observation surface (obtained from the fitted transit time τ_xy_) and average number of molecules within (N_fcs_), from which signal and background contributions were estimated (see section 2.7 for details). (b) Side-view of SLBs in confocal, CH-STED and z-STED modes. Right: axial intensity profiles. (c)-(f) Reconstruction of the effective observation volume and signal to background ratio for confocal (c), 2D STED (d), z-STED (e) and CH STED (f). Intensity profiles (left) were calculated as described in equation 9. Right: axial position-dependent values of total intensity (blue) signal (green) and background (magenta) as determined from equation 10 and as indicated in the legend in panel (f); values represent averages of 2 repeated measurements. The value of SBR integrated across the whole volume is plotted in inserts. STED laser power: 55 mW.

For 2D STED, significant background contributions were detected at distances to the focal plane beyond 300 nm (Figure 4(d)), which is consistent with the results of simulations (Figure 3(b)) and previous work on the subject [9] and corresponds to residual undepleted light from the excitation focus. Smaller relative background values were measured in z-STED (Figure 4(e)) due to the undepleted side lobes that can be readily visualised in imaging (Figure 4(b)), peaking at a distance 700–900 nm from the focal plane. CH-STED (Figure 4(f)) exhibited small background values at depths higher than 300 nm, likely originating from the undepleted area around the optical axis where the intensity of the CH-STED depletion pattern is low. Even though the relative background contribution may appear similarly pronounced in the z-profiles for z- and CH-STED, however, its overall integral relative to signal (SBR values as insets) was considerably lower for CH-STED, as was qualitatively predicted also from the simulations above.

Any quantitative discrepancies between this experiment and simulations likely result from the imperfect model for background noise in simulations (defined as areas with an intensity lower than 1/e^2^ of the peak intensity), which, for instance, is incompatible with the approximation that no background is present at z=0 in experiments. Besides, discrepancies might arise from residual aberrations in the excitation and detection path of our microscope, or from polarisation aberrations that were not modelled in simulations. Nevertheless, both the simulations as well as this depth-profiling experiment nicely illustrated that for all confinement modes background originated from areas of less effective fluorescence depletion far from the focal plane. This was most efficiently supressed in CH-STED, potentiating better STED-FCS performance when measuring 3D diffusion in solution.

## 4 Results

### 4.1 Influence of CH radius parameter

The shape of the CH depletion pattern, and thus of the effective observation volume, can be tuned by varying the size of the inner radius ρ of the CH STED phase mask (Figure 5(a)) [15]. A smaller CH inner radius corresponds to a larger deviation from the vortex pattern, meaning a better axial confinement, at the expense of a lower lateral confinement. To study the impact of CH radius on STED-FCS performance, we measured the diffusion of Abberior Star Red dyes in aqueous solution.

**Figure 5:**
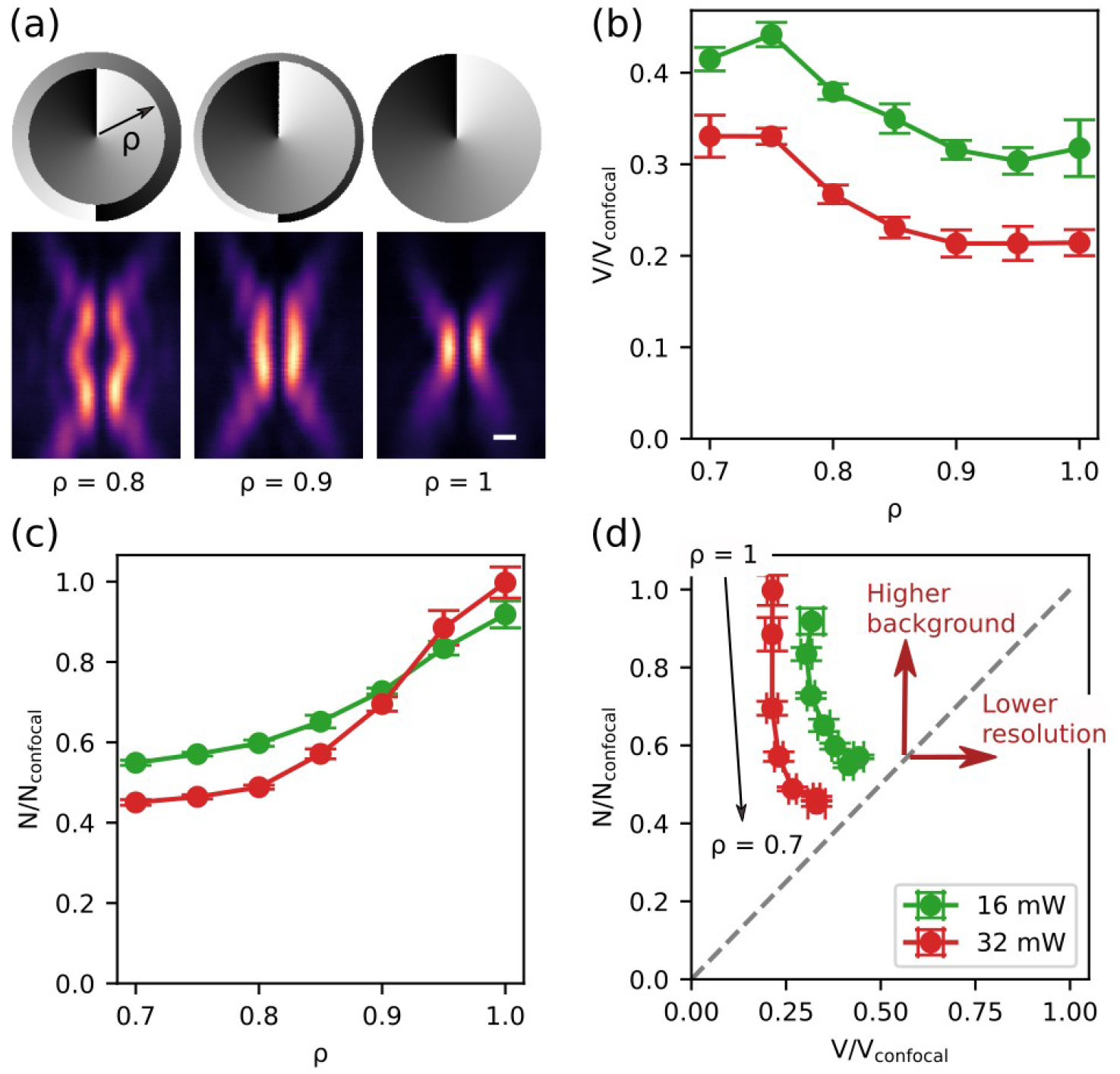
*Influence of CH parameter ρ, as experimentally determined from measurements on an aqueous solution of Abberior Star Red. (a) The inner radius ρ of the central π phase step of the STED phase mask (top) determines the concavity of the effective depletion pattern Bottom: pictures of the xz cross-sections of the depletion pattern imaged using scattering gold beads. Scalebar: 300 nm. (b)-(c) Average size of the effective observation volume (b) and average number of molecules in the observation volume (c) determined from STED-FCS recordings for different values of the parameter ρ and at two different STED laser power, as indicated in the legend in panel (d), and normalised with the confocal value. (d) Variation of observed number of molecules in the observation volume with size of the observation volume, for different values of the parameter* ρ within the range 0.7-1.0 in steps of 0.05. *If no undepleted background is present, the normalised observation volume and number of molecules are equal (gray line). Errorbar: standard deviation, n = 6*.

Aberration correction was performed prior to data acquisition, and the resulting correction was used for all the measurements. We acquired a series of STED-FCS measurements for a range of values of the inner radius parameter, at two different STED laser powers to ensure that any effect observed was not power-specific. Fitting the curves allowed extraction of both the observation volume (calculated from transit times) and of the number of molecules (Figure 5(b)-(c)). Unsurprisingly, STED-FCS recordings with a low CH-STED radius exhibited a lower resolution (Figure 5(b)), but also showed much lower values of the average number of molecules in the observation volume (Figure 5(c)), which indicates a pronounced decrease in background levels. As a measure of the amount of undepleted background, we compared the relative decrease in average number of molecules in the observation volume with that of the observation volume (Figure 5(d)), which were strictly proportional in the absence of any background. This confirmed that lower CH-STED radii depleted the background most effectively. Throughout the rest of this paper, and unless specified otherwise (see section 4.3), we set *ρ* to an intermediate value of 0.85 to benefit from both good resolution and low background levels, which also corresponds to the conditions used previously for CH-STED imaging [15].

### 4.2 Comparison between STED confinement modes in solution

To compare the STED-FCS performance of different confinement modes for characterising 3D diffusion, we conducted STED-FCS experiments in a solution of freely diffusing Abberior Star Red dyes. At a given STED laser power, the resulting FCS curves acquired with CH-STED featured the highest amplitudes (Figure 6(a)), which could either originate from better signal confinement or better noise suppression. Fitting the FCS curves with the model from equation 1 allowed extraction of observation volumes and average number of molecules in the observation volume (Figure 6(b)-(c)). This confirmed that z- and 3D-STED offer the best resolution (Figure 6(b)), but that CH-STED minimises the average number of molecules in the observation volume, suggesting a more efficient background suppression (Figure 6(d)). To verify that background noise levels in CH-STED were in fact lower than they were with other confinement modes, we calculated for each curve the rootmean-square of the fitting residuals normalized to the amplitude (nRMSD) (Figure 6(e)). Lower average residuals again confirmed that CH-STED reduced the background noise with the highest efficiency.

**Figure 6:**
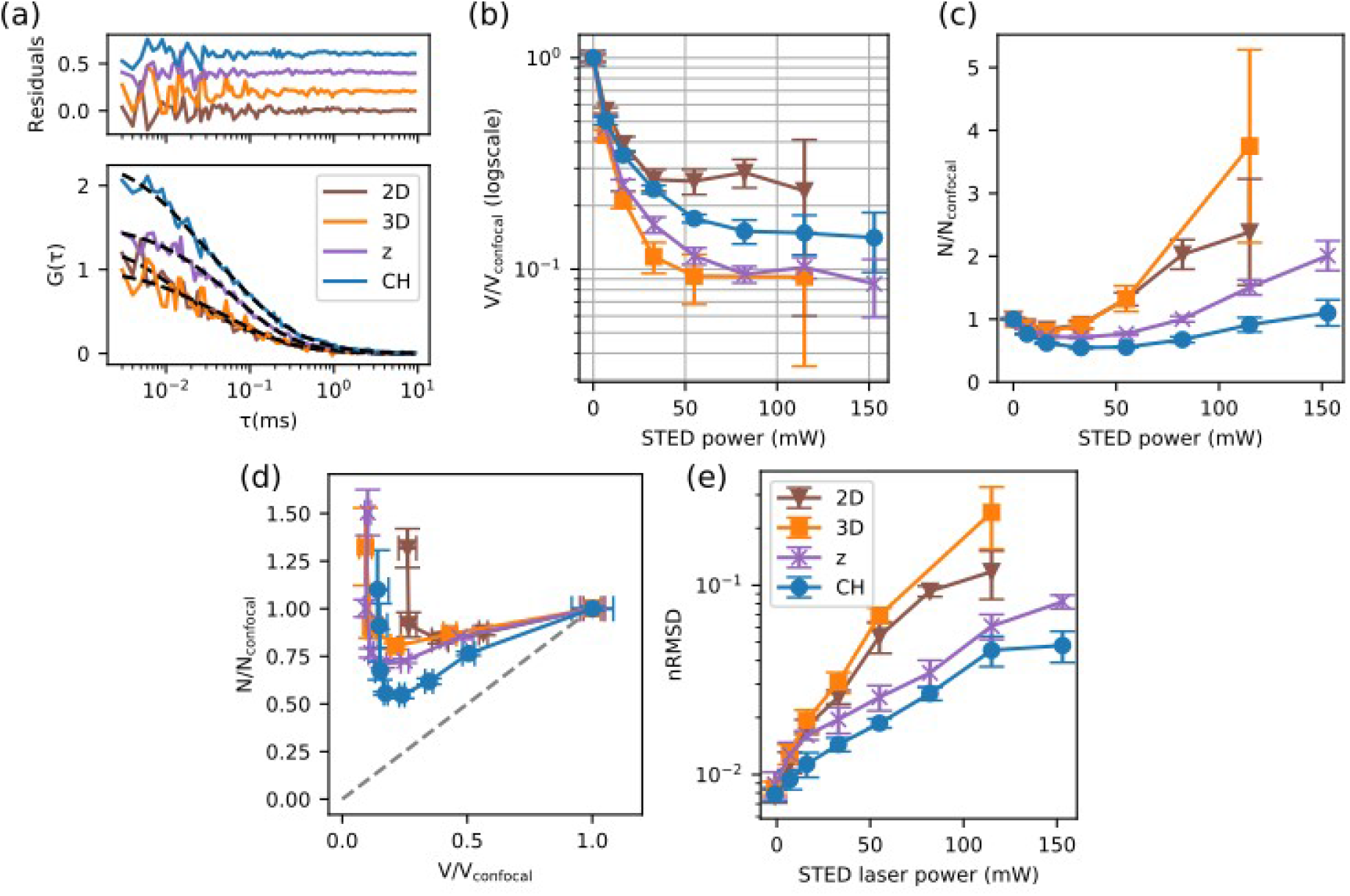
STED-FCS with different confinement modes in an aqueous solution of freely diffusing Abberior Star Red dyes. (a) Representative FCS curves at a STED laser power of 55 mW, for different confinement modes as indicated in the legend. (b-c) STED laser power-dependent effective observation volume (b) and average number of molecules in the observation volume (c), normalised with values determined from confocal recordings and (d) their pairwise scatterplot. STED-FCS recordings without any background would follow the dotted grey line (proportionality between number of molecules in the observation volume and size of the observation volume). (e) Variation of nRMSD (indicator of noise in FCS curves) with STED laser power, for different STED confinement modes as indicated in the legend. Errorbars: s.d, n=6.

### 4.3 Resistance against aberrations

To assess the robustness of STED-FCS measurements with different confinement modes, we used the wavefront shaping capabilities of the SLM to quantify the effects of optical aberrations introduced as low-order Zernike modes, which had previously been reported to be most often present while imaging common biological specimens [25]. We introduced either 0, 0.5 or 1 rad rms of each mode using the SLM, corresponding respectively to no aberrations, to the maximum amplitude of aberrations we experienced in our microscope across this study (see supplementary section S2), and to highly aberrating situations that can be encountered for instance when focussing deep inside a medium with a refractive index mismatch [13], or when focussing deep through an optically inhomogeneous specimen. For each aberration value, we acquired a set of STED-FCS measurement of freely diffusing Abberior Star Red dye in aqueous solution. From the parameters of fits to the obtained FCS curves (Figure 7(a)-(b)), relative variations in the average number of molecules within the observation volume and effective size of the observation volume were used as indicators of the sensitivity of each STED confinement mode to aberrations (Figure 7(c)-(f)). The CH radius was set to a value of 0.75, smaller than in the rest of this study, to maximise the difference between CH STED and 2D STED.

**Figure 7:**
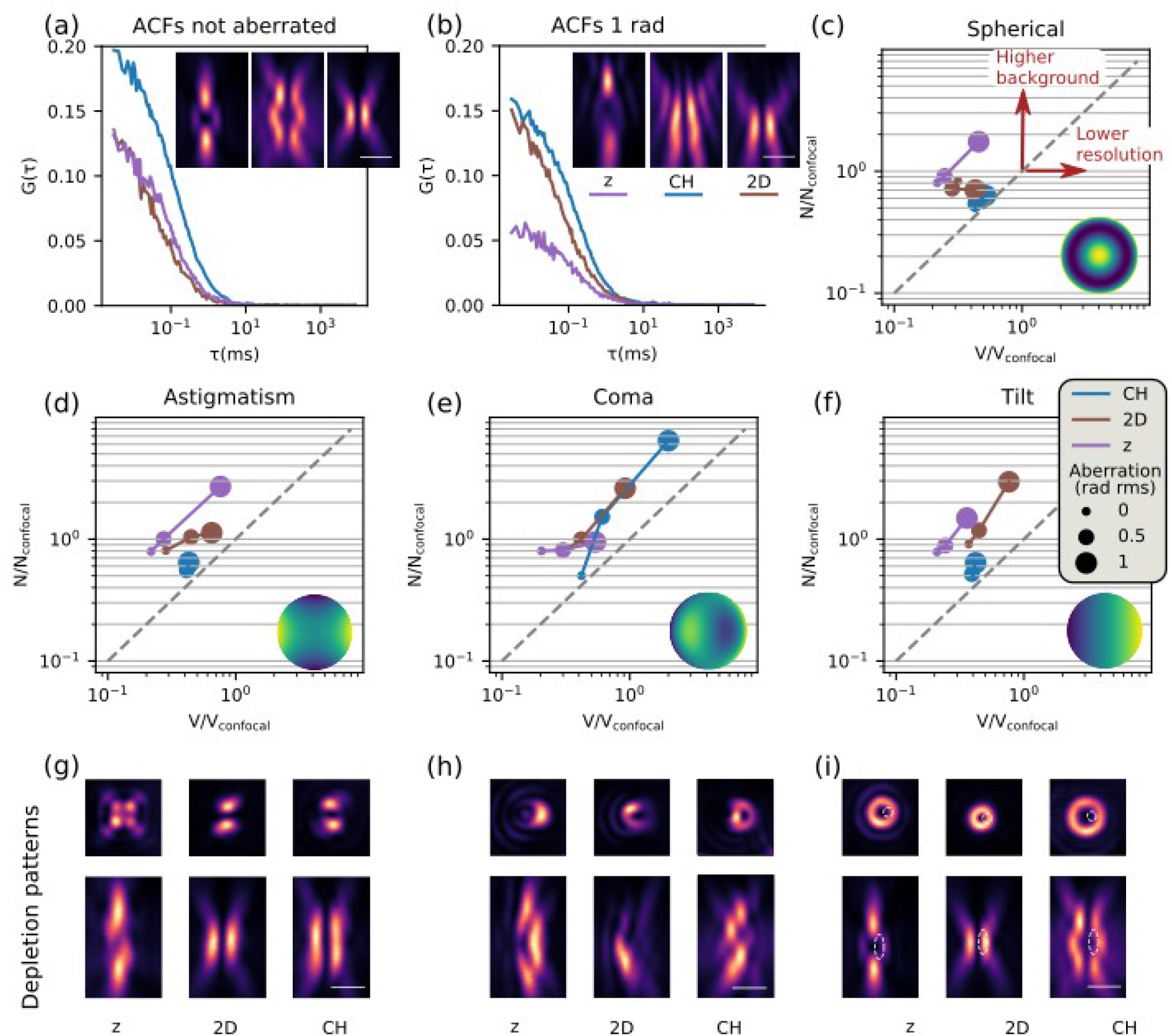
Effect of common aberrations on the depletion pattern on STED-FCS experiments. (a)-(c) Spherical aberration. (a)-(b) FCS curves with different confinement modes, without (b) and with (c) 1 rad rms of spherical aberration introduced in the depletion beam. Images (xz) of the corresponding depletion patterns obtained with a sample of scattering gold beads (insets, scalebar: 1μm). (c) Relative variation of size of the observation volume (x-axis) and average number of molecules in the observation volume (y-axis) upon various amounts of introduced spherical aberration (indicated by the size of the marker) for different STED-FCS confinement modes, as indicated in the legend in panel (f). Aberrations either reduce the resolution or increase the strength of background contributions, as indicated by the arrows. The inset represents the phase distribution of 1 rad rms of spherical aberration. (d)-(i) Corresponding effects of (d), (g) astigmatism (e), (h) coma and (f), (i) tilt on STED-FCS parameters (d)-(f) and gold beads images (g)-(h) in xy (top) and xz (bottom) planes for the depletion patterns with 1 rad rms of the corresponding aberrations. In panel (i), the white dotted line represents the position of the excitation focus. STED laser power: 16 mW.

In accordance with previous work on the effect of aberrations on STED depletion patterns [11], [21], we found that spherical aberrations were detrimental to z-STED but had very limited effects in 2D STED (Figure 7(c)). Suprisingly, spherical aberrations slightly decreased the average number of molecules in the observation volume in 2D STED, which can be attributed to an elongation of the depletion pattern (insets in Figure 7(a)-(b)), leading to a better overlap between excitation and depletion foci. We also found that as predicted in ref. [15], CH-STED was much more resistant to spherical aberrations than z-STED (Figure 7(c)).

Similarly as the spherical aberration, astigmatism (Figure 7(d)) had a more damaging effect on z-STED than on 2D and CH-STED. Coma aberration revealed to be particularly detrimental to CH-STED (Figure 7(e)), as well as for 2D-STED, but not to z-STED. Coma aberration typically occurs when the coverslip is tilted. The exact amount depends on the immersion medium, as more aberrations appear when the index mismatch between the coverslip and the immersion medium is large. For example, previous research suggests that when using a water immersion objective together with a standard, 170 μm thick coverslip, a tilt of 2° of the coverslip would cause approximately 3 rad rms of coma aberrations [26]. In the case of using an oil immersion objective, however, the effect should be much less pronounced. In our system, we never measured coma values larger than 0.2 rad rms (see supplement S2), for which CH-STED would still sport lower background than 2D- and z-STED.

Finally, the impact of tilt, which in this situation corresponds to a misalignment between excitation and depletion foci, is useful to be assessed to estimate the impact of chromatic aberrations [27] or thermal and mechanical drift [13]. We found that 2D-STED was the most sensitive to tilt, followed by z-STED and CH-STED, which can be linked to the respective sizes of their central areas, with a smaller central area leading to a higher vulnerability to misalignment (Figure 7(f)).

To help illustrate these results, we defined the degree of sensitivity to aberrations as follows: a given STED depletion pattern is defined to have a high sensitivity to a given aberration mode if 0.5 rad rms of this mode increases either the number of molecules in the observation volume or the size of the observation volume by more than 50%. If instead 1 rad rms of this mode leads to such an increase, the sensitivity to this mode is defined as intermediate. Otherwise, the sensitivity is low. Results are presented in Table 1.

**Table 1:** Sensitivity of STED confinement modes to common optical aberrations

|  | Spherical | Astigmatism | Coma | Tilt |
| --- | --- | --- | --- | --- |
| 2D | low | high | intermediate | intermediate |
| CH | low | low | high | low |
| z | intermediate | intermediate | low | intermediate |

### 4.4 STED-FCS in living cells

To evaluate the applicability of each STED confinement mode to measurements of 3D diffusion in biological specimens, we measured 3D diffusion of Silicon-Rhodamine dye in the cytoplasm of human fibroblasts (Figure 8(a)) using z-, 3D- and CH-STED-FCS. 2D-STED was not assessed, as 2D-STED-FCS results already showed much poorer signal and spatial resolution in solution than other confinement modes (see Figure 6). In our experiments, we found that the decrease in observation volume was similar for all three confinement modes (Figure 8(b)), unlike in solution where, at these STED laser powers, 3D-STED exhibited a higher resolution (Figure 6(b)). The difference may be explained by the high variability of measurements in cells. To estimate noise levels, we calculated the nRMSD of each curve (Figure 8(c)). Noise levels were the lowest for CH-STED, followed closely by z-STED FCS, and the highest when using 3D-STED, consistently with measurements in solution (Figure 6(d)-(e)). Note that we could not reliably assess the average number of molecules in the observation volume due to too large cell to cell variations.

**Figure 8:**
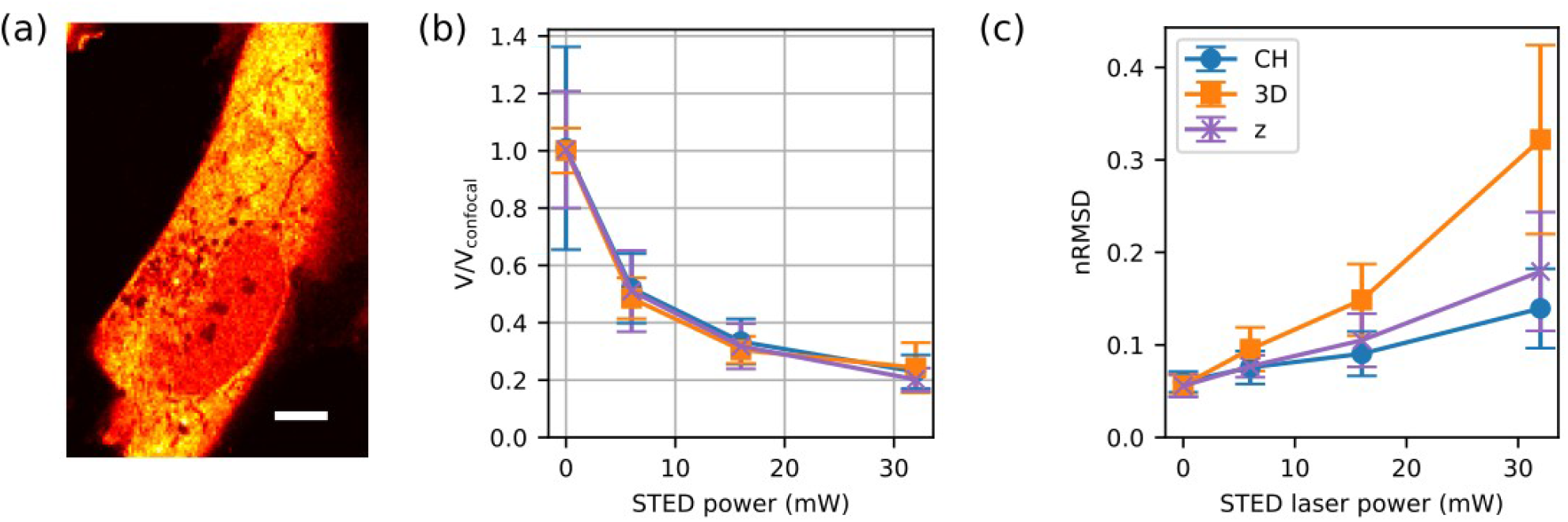
Diffusion in the cytoplasm of living cells recorded with STED-FCS and different confinement modes using the fluorescent and membrane permeable dye Silicon Rhodamine. (a) Confocal xy image of a cell where STED-FCS experiments were performed. (scalebar: 5 μm). (b) Observation volumes at different STED laser powers and with different STED patterns as indicated in the legend in panel (c), normalised with the confocal value, and (c) noise in correlation curves, measured as root-mean-square of the fitting residuals normalized to the amplitude (nRMSD) for different confinement modes, as a function of STED laser power (mean +/− s.d, n>=12 curves from 3 cells).

## 5 Discussion

We investigated the effects of four different STED confinement modes (2D-, z-, 3D- and CH-STED) on the performance of STED-FCS experiments, both theoretically using simulations and experimentally in a variety of systems. Our study shows that background from undepleted areas can significantly deteriorate signal levels in STED-FCS, and this effect largely depends on the depletion pattern used. We found that undepleted background noise was particularly high in 2D- and 3D-STED, while it was reduced with z-STED and minimal with CH-STED. We also found that CH-STED was generally less sensitive to optical aberrations than z-STED, especially to spherical aberrations, which are common in biological experiments. As a result, CH-STED can be considered a depletion pattern of choice for STED-FCS experiments of 3D diffusion, as long as they do not require the highest attainable resolution. In this case, z-STED combined with aberration correction should be preferred.

We showed that CH-STED is a tool of choice to facilitate STED-FCS experiments in 3D environments with STED setups incorporating a SLM. The new understanding of the origin of background in STED-FCS will also help designing new depletion patterns that optimise background reduction. In STED systems with a double-pass configuration, for example, we expect a combination of z- and CH-STED to join the great resolution of the former and the background suppression capabilities of the latter for optimised STED-FCS in 3D.

## Acknowledgements

The authors thank Dr Katharina Reglinski for providing us with plasmids, and Dr Erdinc Sezgin for his help regarding sample preparation.

## Data availability

The research materials supporting this publication can be accessed by contacting the authors.

