## Supplementary text for "Background reduction in STED-FCS using coherent-hybrid STED"

### Supplement

#### 1 Data fitting

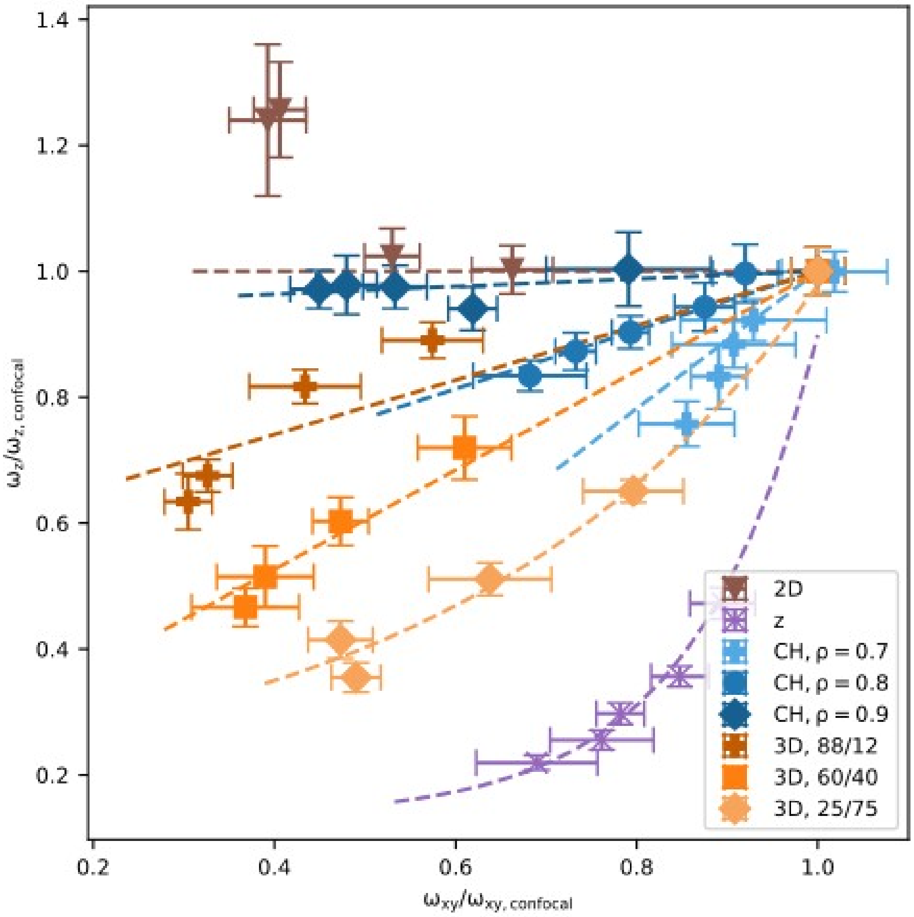


*Figure 1: Measurement of resolution variations with STED laser power using fluorescent beads, with the oil immersion objective for different STED confinment modes, plotted as two-dimensional plot of value pairs of lateral $\omega_{xy}$ and axial $\omega_z$ $1/e^2$ radius, normalised with confocal values.(mean +/- std, n = 10 beads per datapoint).*

We designed the fitting methods as described in [1]. Because conventional fluorophores do not have a high enough molecular brightness to fit at the same time axial and lateral transit times, we instead fitted the observation volume with a prescribed shape. The exact shape of the observation volume depends on the STED confinement mode and was determined from pictures of immobilised 40 nm crimson beads.

For each STED confinement mode and at STED laser powers ranging from 0 (confocal) to 150 mW we acquired a set of three-dimensional image stacks of fluorescent beads. In each of the stacks, 7 to 10 beads were manually selected, and their axial cross-section was fitted with a two-dimensional gaussian ellipsoid to estimate their lateral and axial full width at half maximum. Results are presented in Figure 1. The variation of axial resolution with lateral resolution was then fitted to either an exponential or a linear function, allowing description of the shape observation volume with a single parameter instead of two.

### 2 Aberrations

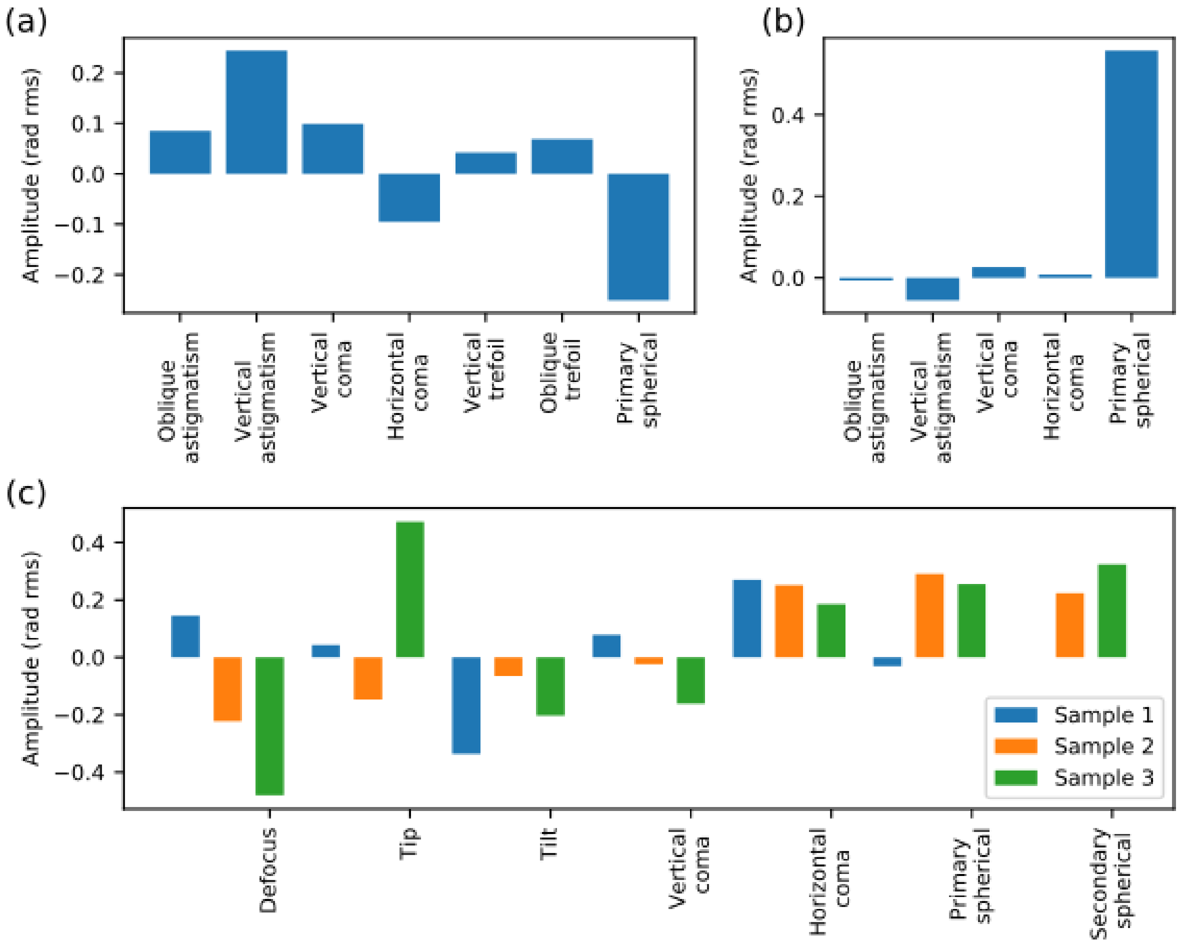


*Figure 2: Aberration values measured experimentally. (a) System-induced aberrations, (b) typical aberration values found when measuring free diffusion in a solution of Abberior Star Red dyes and (c) aberrations measured in three cells samples, as indicated in the legend.*

Aberrations in the depletion beam were corrected to ensure an optimal measurement quality. System-induced aberrations caused by misalignments or imperfections in the optical components were corrected first, using the sensorless method. A  a sample of scattering gold beads was scanned through the focus while various amounts of aberrations were induced by the SLM, and the image quality was assessed using the standard deviation of an image. The amplitude of system aberrations determined with this method was relatively low, limited to around 0.2 radians root mean square (rad rms) (Figure 2(a)). When measuring STED-FCS in a freely diffusing solution of Abberior Star Red dyes at penetration depths comprised between 3 and 5 µm, the refractive index mismatch between the immersion oil of the objective and the solution induced mostly spherical aberrations (Figure 2(b)), with an amplitude approximately equal to 0.5 rad rms. In cells, aberrations measured were more heterogeneous, with maximum amplitudes equal to 0.4 rad rms (Figure 2(c)). Misalignements between excitation and depletion beam occurred because of the effect of thermal and mechanical drift, as we observed previously [1], and were corrected using tip, tilt and defocus.
